# Neural and non-neural contributions to sexual dimorphism of mid-day sleep in Drosophila: A pilot study

**DOI:** 10.1101/027847

**Authors:** M. Khericha, J.B. Kolenchery, E. Tauber

## Abstract

Many of the characteristics associated with mammalian sleep are also observed in *Drosophila*, making the fruit-fly a powerful model organism for studying the genetics of this important process. Among these similarities is the presence of sexual dimorphic sleep patterns, which in flies, is manifested as increased mid-day sleep (‘siesta’) in males, compared to females. Here, we have used targeted miss-expression of the gene *transformer* (*tra*) and *tra2* to either feminise or masculinise specific neural and non-neural tissues in the fly. Feminization of males using three different GAL4 drivers which are expressed in the mushroom bodies induced a female-like reduced siesta, while the masculinisation of females using these drivers triggered the male-like increased siesta. We also observed a similar reversal of sex-specific sleep by miss-expressing *tra* in the fat body, a key tissue in energy metabolism and hormone secretion. In addition, the daily expression levels of *takeout*, an important circadian clock output gene, were sexually dimorphic. Taken together, our experiments suggest that sleep-sexual dimorphism in *Drosophila* is driven by multiple neural and non-neural circuits, within and outside the brain.

## Introduction

Studies in various organisms have shown that various sleep properties are gender specific. In humans for example, the frequency of sleep spindles (a burst of oscillatory neural activity during stage N2 sleep) is elevated in women compared with men (Gaillard, Blois 1981). In addition, women sleep longer, when deprived from external cues under lab conditions (Wever 1984), and slow wave sleep (SWS) is more frequent in women than in men (Reynolds et al., 1990). Sex difference in sleep patterns is also present in mice (Sinton et al., 1981; Paul et al., 2006) and rats (Fang, Fishbein 1996).

Similar to mammals, the pattern of sleep in *Drosophila* is also sexually dimorphic, with a pronounced mid-day sleep (’siesta’) in males, but not in females (Andretic, Shaw 2005; Ho, Sehgal 2005). In addition, the fly response to sleep deprivation has also been studied (Hendricks et al., 2003; Shaw et al., 2002), although gender dimorphic differences have been observed only in the circadian clock mutant cycle (aka *Bmal1*). Female mutants have a pronounced rest rebound, whereas in males the homeostatic response is reduced or non-existing.

A recent study (Catterson et al., 2010) has shown that diet has a major impact on sleep patterns, in a way which was also sex-dependant. Males fed with dietary yeast extracts showed increased locomotor activity and shorten diurnal and nocturnal sleep, while females responded to this diet with reduced daytime locomotor activity and a more fragmented nocturnal sleep. The reduced mid-day sleep in females has been associated mainly with inseminated females (Isaac et al., 2010), which has led to the suggestion that the sex-peptide, a male seminal peptide transferred during copulation, modulates the female behaviour and promotes their mid-day waking.

Sex determination in *Drosophila* has been extensively studied (Schutt, Nothiger 2000) and genetic tools are available, allowing manipulation of specific target tissues. The *transformer* (*tra*) gene is a key gene in the cascade responsible for somatic sexual differentiation. In females, splicing of *tra* (mediated by SXL) generates TRA protein that activates the female sexual differentiation. In males, the *tra* pre-mRNA is spliced into its male-specific form, which translates into a truncated inactive protein, consequently leading to male sexual differentiation. Ectopic expression of the female form of *tra* RNA causes chromosomal males to develop as females (McKeown et al., 1988). The UAS-GAL4 binary system in Drosophila (Brand, Perrimon 1993) allows the expression of the female spliced form of *tra* in targeted cells in a male, inducing a female pattern of development; strains with a GAL4 transgene expressed in a defined set of cells are crossed to those carrying the female-specific *tra^F^* fused to upstream activating sequence (UAS-*tra*). This leads to activation of *tra* in all the tissues expressing GAL4, creating tissue-specific feminization (Ferveur et al., 1995; Ferveur et al., 1997). A similar approach has also been used to masculinise female specific tissues, using a *tra*-2 RNA interfering construct (UAS-*tra2*-IR) (Lazareva et al., 2007). Here, we have used the UAS-GAL4 system to feminize male specific regions of the brain and masculinise female specific neurons, trying to identify the sleep circuits that may be controlling this sexually dimorphic behaviour in flies.

## Materials and Methods

### Fly strains

To feminise males, the strain w; UAS-*tra^F^* from the Bloomington Drosophila Stock Centre at Indiana University (stock number 4590) was used. For female masculinisation, we used a transgenic strain carrying dsRNAi construct targeting *tra2*, (UAS-*tra2*-IR), which was obtained from Vienna Drosophila RNAi Centre (stock v8868). Another strain targeting UAS-*tra* also has been used (stock v2560), but preliminary tests indicated that mid-day sleep females UAS-tra-IR is unusually high, and therefore not useful for testing female masculinisation. UAS-*dicer2* transgenic strain (stock v60008) was used to enhance the efficiency of RNAi in some crosses (specified when used).

Four GAL4 enhancer-trap strains, 103Y, 30Y, 121Y (Gatti et al., 2000) and Voila-GAL4 (Balakireva et al., 1998) driving expression in the mushroom bodies (MB), central complex and a small cluster in pars intercerebralis (PI) were a gift from Jean-François Ferveur at the University of Dijon. Additional GAL4 strains were obtained from Bloomington Stock Centre included the pan neural *w;elav-GAL4* (stock 8760), and w;1471-GAL4 strain with expression patterns in the γ lobes of MB (stock 9465). *takeout*-GAL4 driving expression in the fat body as well as in a subset of cells within the maxillary palps and antennae (Dauwalder et al., 2002) was a gift from Brigitte Dauwalder at the University of Houston.

Each of the strains above was also crossed to *w^1118^* and their F1 progeny were used as two controls (UAS and GAL4) compared to the phenotype of flies carrying the both transgenes. All stocks and experimental crosses were maintained at 25°C with a Light:Dark 12:12 cycle, and kept on standard cornmeal/sugar-based food.

### Sleep assay

The sleep/wake pattern of flies aged 3-4 days was monitored using the Drosophila Activity Monitoring System (DAMS, TriKinetics) at 25°C in LD 12:12, for a total of 4 days. Only virgin females were used in all experiments. Data were collected in five-min bins, and sleep was quantified by summing consecutive bins for which no activity was recorded, using the R software (R Development Core Team 2010). Since the mid-day ’Siesta’ sleep time interval varied among strains (typically, between 5-8hr after light on), we quantified the average mid-day sleep during two hours around noon (5-7 hr after lights on). This has simplified the algorithm and ensured the capture of mid-day sleep. In the feminizing experiments, where the female-spliced form of *tra* was expressed in males, siesta sleep was calculated both in the feminized males and in females, and compared to their background controls. Similarly, in masculinisation of the females, RNAi constructs of *tra* and tra-2 were expressed in females, and siesta sleep was assessed in males and masculinised females, and compared to their background controls. In each experiment, the sleep scores of the three genotypes were compared by Kruskal-Wallis ANOVA. Tests indicating significant difference were followed by a non-parametric post-hoc test (Siegel 1988) (p.213-214), comparing each of the control to the GAL4/UAS genotype. Statistical tests were carried with the *pgirmess* library implemented in the statistical software “R” (R Development Core Team 2010).

### RNA Quantification

The mRNA levels of *takeout* were assayed by qPCR. We analysed males, virgin females and mated females, 4-5 days old. Flies were maintained at 25°C in a 12-h LD cycle for 5 days. On the sixth day the files were collected at two different time points, immediately after lights on (Zt0), and six hours after lights off (Zt6). Total RNA was isolated from male fly heads using TRIZOL (Invitrogen). 500 ng of total RNA was used for cDNA synthesis, which was carried with the Affinity Script kit (Stratagene). OligodT primers were used for the first strand synthesis. qPCR was carried using a SYBR Green assay (Agilent technology). The standard curve method was followed to quantify *takeout* mRNA, in 25 μL reactions, with 0.3 μM of final primer concentration. The forward primer was, 5’-GCCTTTTGGTCTCGGTGGAT-3’; reverse primer, 5’-TCCCCATTCTTCACCAGCG (amplicon size 142bp). *Ribosomal protein 49* mRNA (*rp49*) was used as the reference gene. The forward primer was, 5’-TTACAAGGAGACGGCCAAAG the reverse primer: 5’-CTCTGCCCACTTGAAGAGC.

## Results and Discussion

All the transgenic strains used in this study exhibited a marked sexual dimorphic in mid-day sleep (Figures S1-S5 in Supplementary Material), with males sleeping up to twice as much as females (males [mean±SD]: 94 ± 21, females: 42 ± 24 min during 2 hr at mid-day), similar to the previously reported sleep differences exhibited by wild-type Canton-S (Andretic, Shaw 2005; Ho, Sehgal 2005).

The mushroom bodies (MBs) have been previously implicated as a key brain structure in sleep regulation (Joiner et al., 2006; Pitman et al., 2006). The role of the MB seems to be complex: preventing the MB output (either transiently, or by ablation) results in reduced sleep (Joiner et al., 2006; Pitman et al., 2006), but raising the activity of Go signalling in the MB enhances sleep (Guo et al., 2011). This complexity has been evident in a recent study showing that Go signalling is present in two adjacent subtypes of MB cholinergic neurons that play opposite roles in sleep regulation (Yi et al., 2013). Most parts of the MB are innervated by a single pair of neurons, the dorsal paired medial (DPM), which have recently shown to promote sleep (Haynes et al., 2015). The mechanism involves inhibition of the MB α’/β’ neurons, by GABA release. The MB outputs converge onto a small subset of neurons (called MB output neurons, MBONs), whose role in sleep regulation has been recently studied in detail (Aso et al., 2014b). Glutamatergic MBONs were found to be sleep-suppressing while GABAergic or cholinergic neurons were sleep-promoting.

Given the role of MB in sleep, we have tested the contribution of these brain structures to sexual dimorphic sleep. The 121Y-GAL4 strain drives expression in the central complex (CC) and the MBs (Gatti et al., 2000; Armstrong et al., 1995). Using this driver for expressing UAS-*tra* (Figure 1A) resulted in significantly reduced (feminised) male siesta sleep compared to control males carrying only a single transgene. Using this driver for knocking down *tra2* for masculinisation of the MBs induced siesta sleep in females, which was significantly higher than either of the single transgene controls (Figure 1B). Note that the similar sleep level in females in the feminisation experiment (Figure 1A) or the males in the masculinisation experiment (Figure 1B) suggest that the response that we observe is not merely due to the interaction between the GAL4 and UAS genetic backgrounds.

**Fig. 1.**
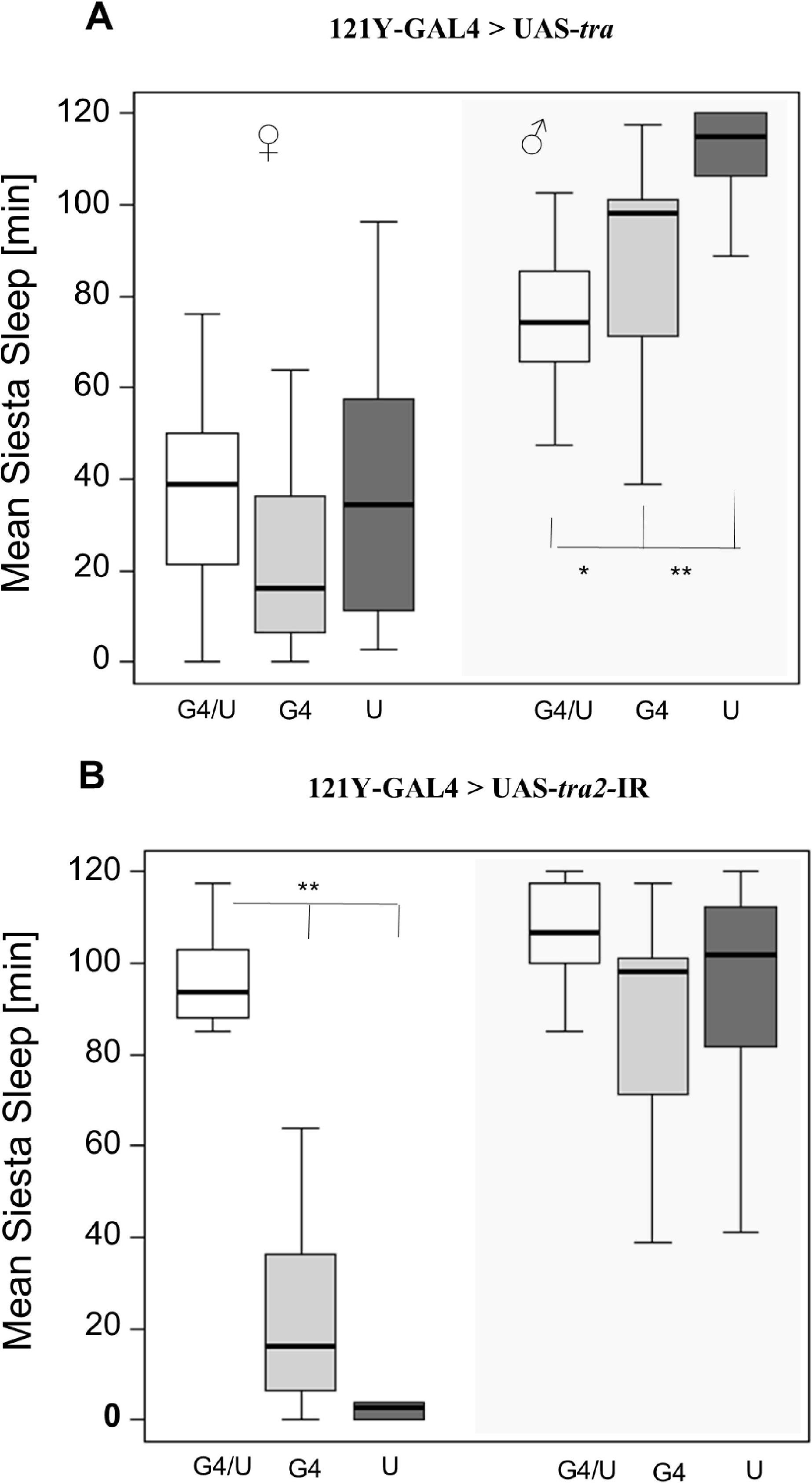
Siesta sleep following feminsation and masculinisation of MBs. Box-plots showing siesta sleep in flies carrying 121Y-GAL4 driving **(A)** UAS-tra (feminsation of males), and **(B)** UAS-tra2 (masculinisation of females). In each panel, the three left boxes show sleep in females, and the three right boxes (shaded grey) are for the males. The data shown in each panel represent siesta sleep for the GAL4/UAS genotypes (white, n= > 20 for all GAL4 lines; males and females) and the single transgene control genotypes (GAL4/+; light grey, UAS/+; dark grey) for both sexes. Asterisks represent experimental genotype (GAL4/UAS) significance levels compared to control genotypes (GAL4/+ and UAS/+). Non-parametric post hoc tests were performed (*P < 0.05, **P < 0.01). The line within each box represents the median siesta sleep averaged over 4 days (in minutes), and the boxes extend to 25 and 75 percentiles. Note that significance differences are only tested for males in the feminisation experiments, or females in the masculinisation experiments.

The 30Y-GAL4 transgene is expressed in the MBs and the CC (Gatti et al., 2000; Yang et al., 1995). Feminisation of males using this driver induced a small, but significant, reduction of sleep compared to the UAS control (p < 0.01), but not compared to the GAL4 control, which showed unusual reduced sleep (Figure 2A). The effect of using this driver to masculinise females was stronger, and UAS-tra2-IR (Figure 2B) resulted in siesta sleep in females that was comparable to that exhibited by males.

**Fig. 2.**
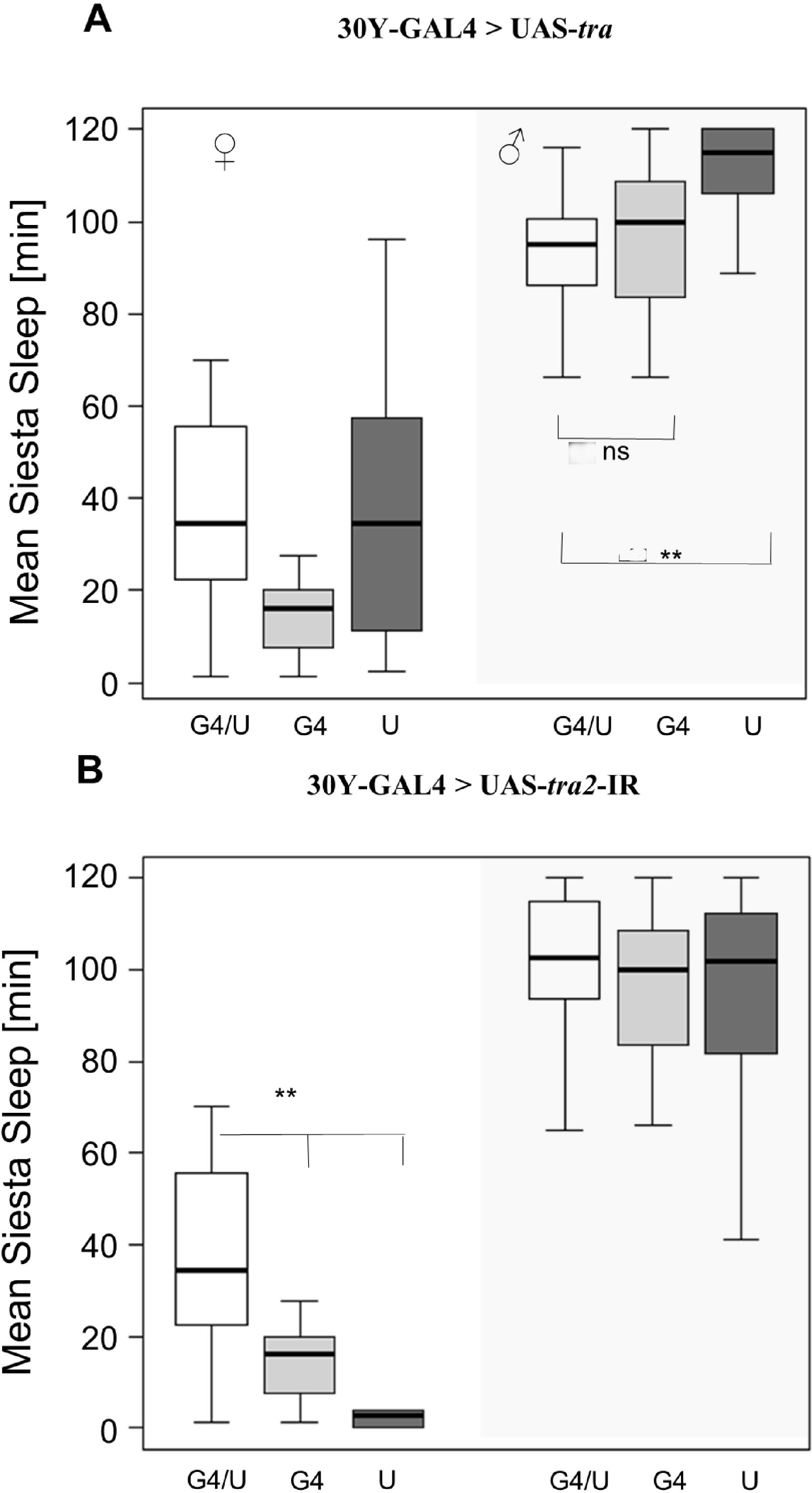
Feminsation and masculinisation of MBs using 30Y-GAL4. Box-plots showing siesta sleep in flies carrying 30Y-GAL4 driving **(A)** UAS-*tra* (feminsation of males), and **(B)** UAS-tra2 (masculinisation of females).Plotting scheme same as in Fig. 1.

Using the 103Y-GAL4 line whose expression also extend to the MBs and CC (Tettamanti et al., 1997) also induced reversal of siesta sleep; in males, sleep was reduced compared to the UAS control (but not compared with the GAL4 control, which showed non-typical low siesta, Figure 3A). In females, brain masculinisation induced male-like siesta sleep (Figure 3B). We observed a similar reversal of sleep using the 1471-GAL4 which is expressed in the γ lobes of MBs (Isabel et al., 2004) (Figure 4). In contrast, using the *Voila*-GAL4 line, which is expressed in the MBs and the antennal lobes (Balakireva et al., 1998), did not result in any significant change in sleep in either feminised males or masculinised females (Figure S6 in Supplementary Material).

**Fig. 3.**
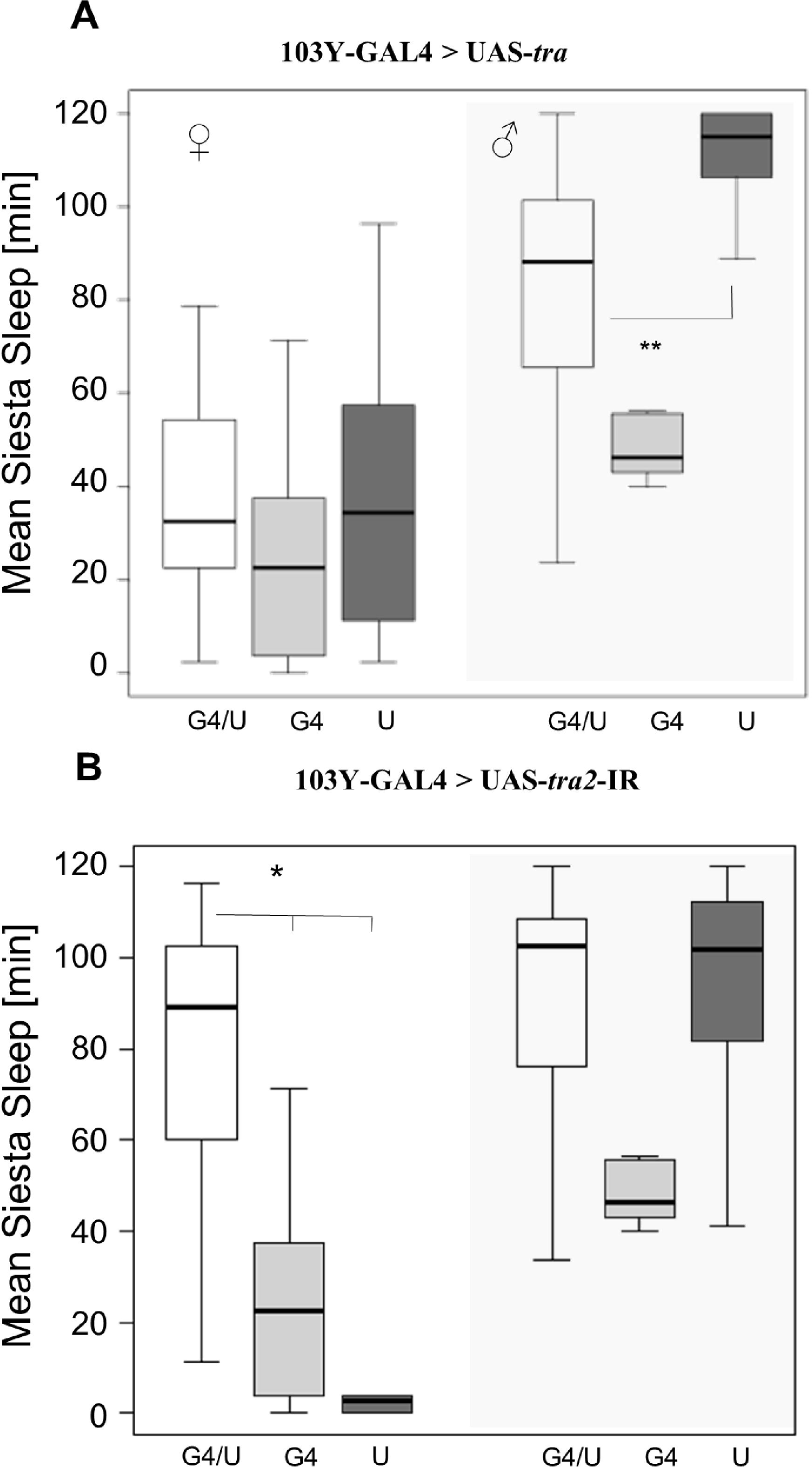
Feminsation and masculinisation of MBs using 103Y-GAL4. Box-plots showing siesta sleep in flies carrying **103Y-GAL4** driving **(A)** UAS-tra (feminsation of males), and **(B)** UAS-tra2 (masculinisation of females).Plotting scheme same as in Fig. 1.

Four of the abovementioned lines 121Y, 30Y, 103Y and Voila have been previously implicated in controlling a sexually dimorphic locomotion behaviour (Gatti et al., 2000), with males exhibiting significantly shorter inter-bout intervals (and lower variation) than females. The overlap of the expression patterns of these GAL4 lines was restricted to a small cluster in the pars-intercerbralis (PI), which was therefore suggested as a candidate for the location of that circuit. Here, however, the Voila driver did not have any effect on reversing sleep, while the driver *1471*-GAL4 (not expressed in the PI) did (Figure 4). Given that the overlap between these driver lines mainly consists of the MBs, which have recently been implicated in the regulation of sleep (Joiner et al., 2006; Pitman et al., 2006), it is likely that neurons in this centre also underlie the variations in siesta sleep. We do note however that in three of our feminisation experiments (Figure 2A, 3A, 4A) the experimental line did not differ significantly from the GAL4 driver, complicating our interpretation. Interestingly, males carrying these MB GAL4 driver showed unusual sleep, which may be the result of GAL4 accumulation in brain neurons as was previously reported (Rezaval et al., 2007). Testing additional GAL4 drivers with more specific expression in the MBs, for example by using the split-GAL4 collection that has been recently created (Aso et al., 2014a), will aid identifying the neurons underlying sexual dimorphism. In addition, given that the PI has been shown to be important for sleep regulation (Foltenyi et al., 2007; Crocker et al., 2010) further analysis using PI-specific drivers would help ruling out a role for this brain region in the sexual dimorphism. Future experiments would also benefit from backcrossing all GAL4 and UAS strains onto a uniform genetic background, which is rather important in sleep studies involving genetic screens (Axelrod et al., 2015).

**Fig. 4.**
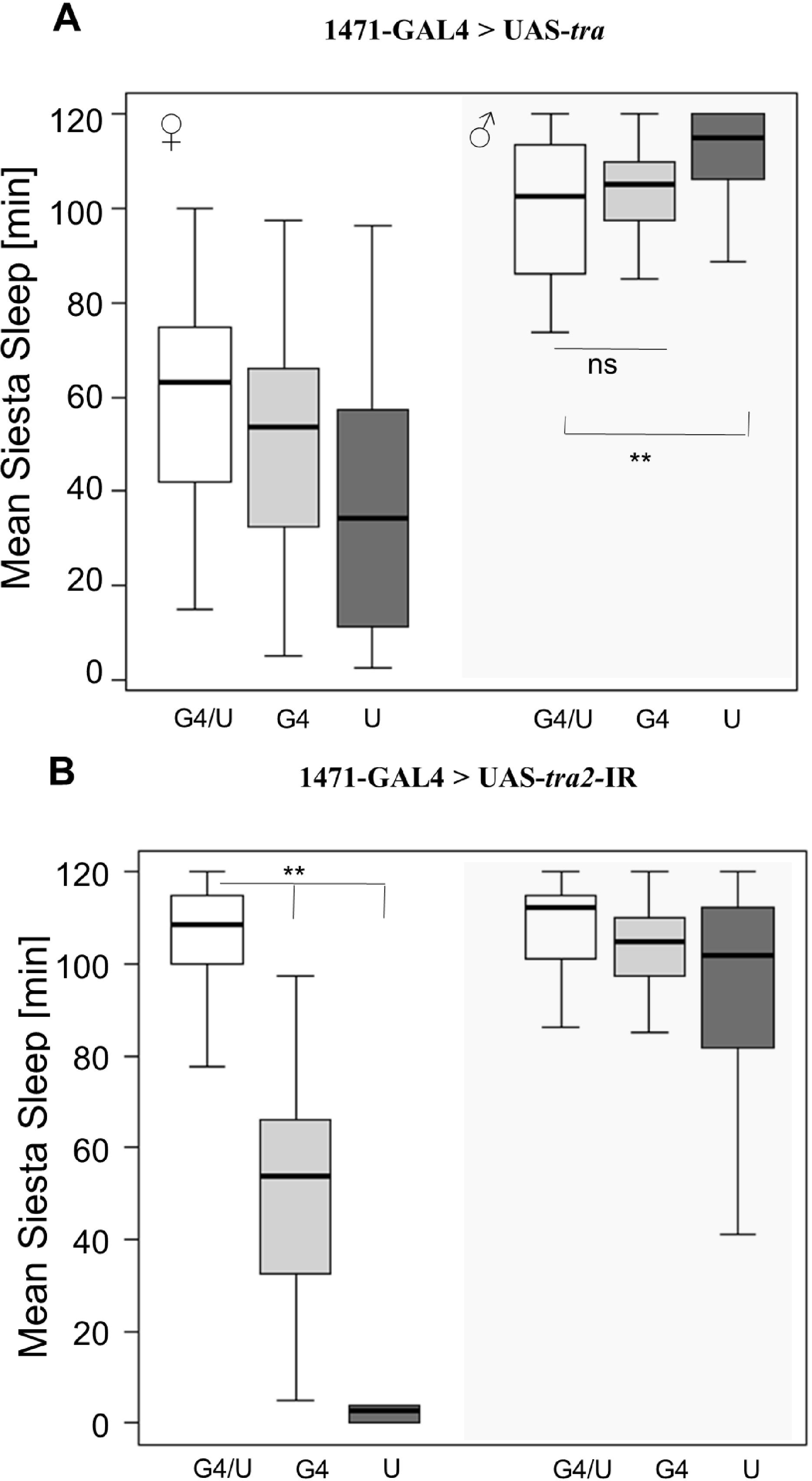
Feminsation and masculinisation of MBs using 103Y-GAL4. Box-plots showing siesta sleep in flies carrying **1471-GAL4** driving **(A)** UAS-tra (feminsation of males), and **(B)** UAS-tra2 (masculinisation of females).Plotting scheme same as in Fig. 1.

The use of GAL4 lines may be combined with the GAL80 enhancer traps, to repress the GAL4 expression, to drive feminization or masculinisation in a subset of cells of the drivers described here, refining the candidate regions (Suster et al., 2004). This approach has been very successful in refining the brain neurons that constitute the circadian clock in *Drosophila* (Stoleru et al., 2004).

Interestingly, the *takeout*-GAL4 strain, which is expressed in the fat body (Dauwalder et al., 2002) was also effective in reversing sleep (Figure 5). While feminisation of males caused only small reduction of siesta sleep (compared with the UAS, but not with the GAL4 control), the masculinisation of females using the UAS-tra2-IR transgene induced a substantial increase in siesta sleep in females (Figure 5), indicating a role for the fat body in sleep sexual dimorphism. Previous studies showed that *takeout* is under circadian control (Benito et al., 2010) and is involved in the regulation of feeding as well as adaptation to starvation (Meunier et al., 2007; Sarov-Blat et al., 2000). Thus, it is possible that the sleep sexual dimorphism is mediated by *to* (and the fat body) indirectly, so feminizing or masculinising the fat body changes the feeding status of the animal, and consequently its foraging behaviour. This idea fits well the recent studies that show a direct link between sleep pattern and feeding (Catterson et al., 2010).

**Fig. 5.**
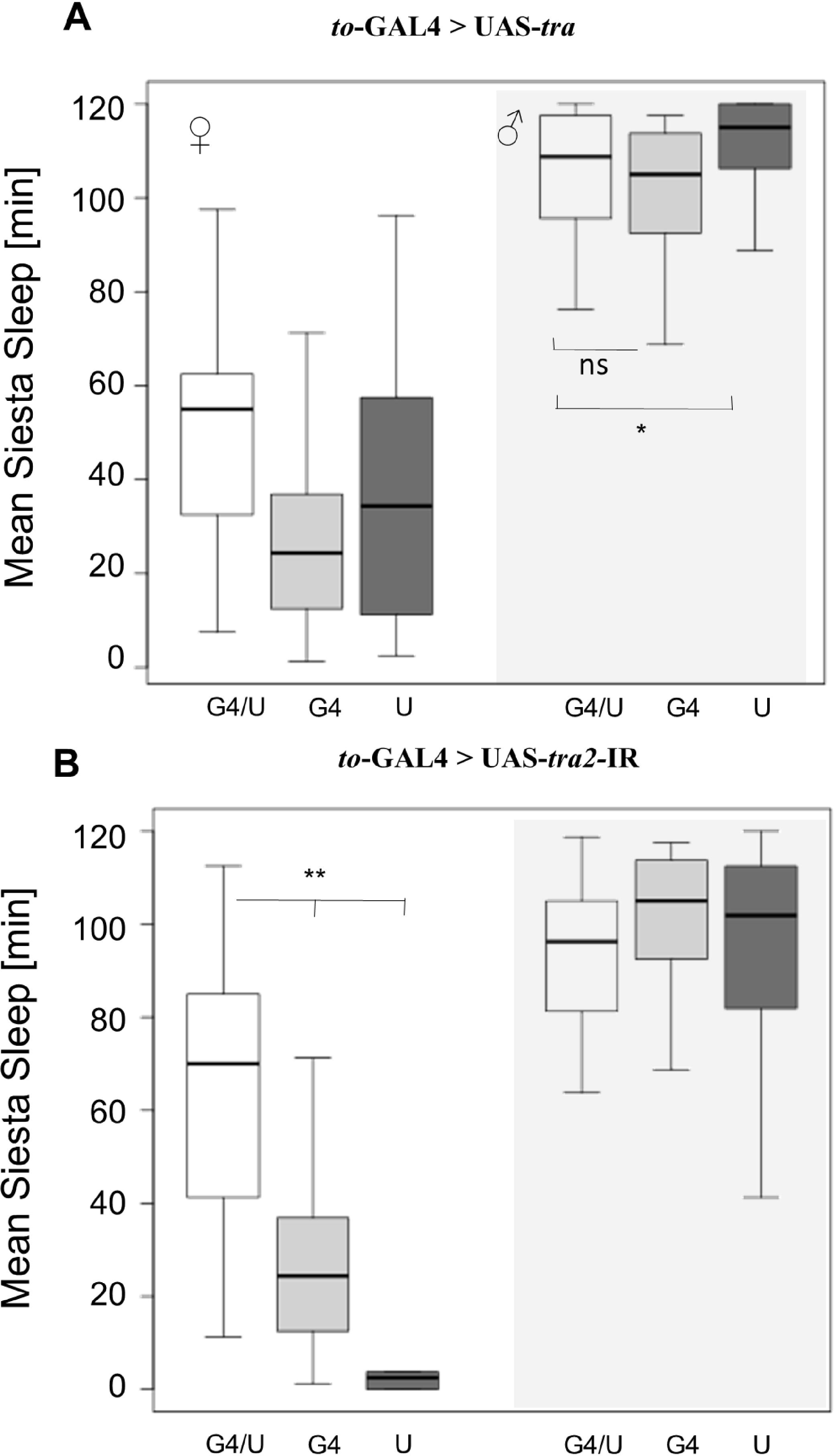
Siesta sleep following feminisation and masculinisation of the fat body. The *takeout (to)* Gal4 driver was used for **(A)** feminsation of males using UAS-tra^F^, and **(B)** masculinisation of females using UAS-traz-IR. Plot parameters are as described in Figure 1.

We have also analysed the transcript level of *to* during the beginning of the day (Zt0) and midday (Zt6) (Figure 6). The expression of *to* was sexually dimorphic with a significant time-sex interaction (F_1,10_=4.99, p < 0.05). In both males and females, transcript level was relatively high at the beginning of the day and decline at midday as was previously reported (Benito et al., 2010), but was substantially higher in males at Zt0 (Figure 6). Thus, sex-dependent differences in *to* expression at the beginning of the day may contribute to the differences in siesta sleep. Although RNA level converged to the same level at midday in males and females, there might be a time-lag between the mRNA and the protein profiles, leading to different TO protein level between males and female just before siesta time (although previous studies suggested that this lag is rather small, (So et al., 2000; Benito et al., 2010). Interestingly, in a recent study that analysed sleep behaviour in wild populations over a broad latitudinal range (Svetec et al., 2015), *to* was identified as a strongly differentially expressed gene, suggesting that it is the target for natural selection.

**Fig. 6.**
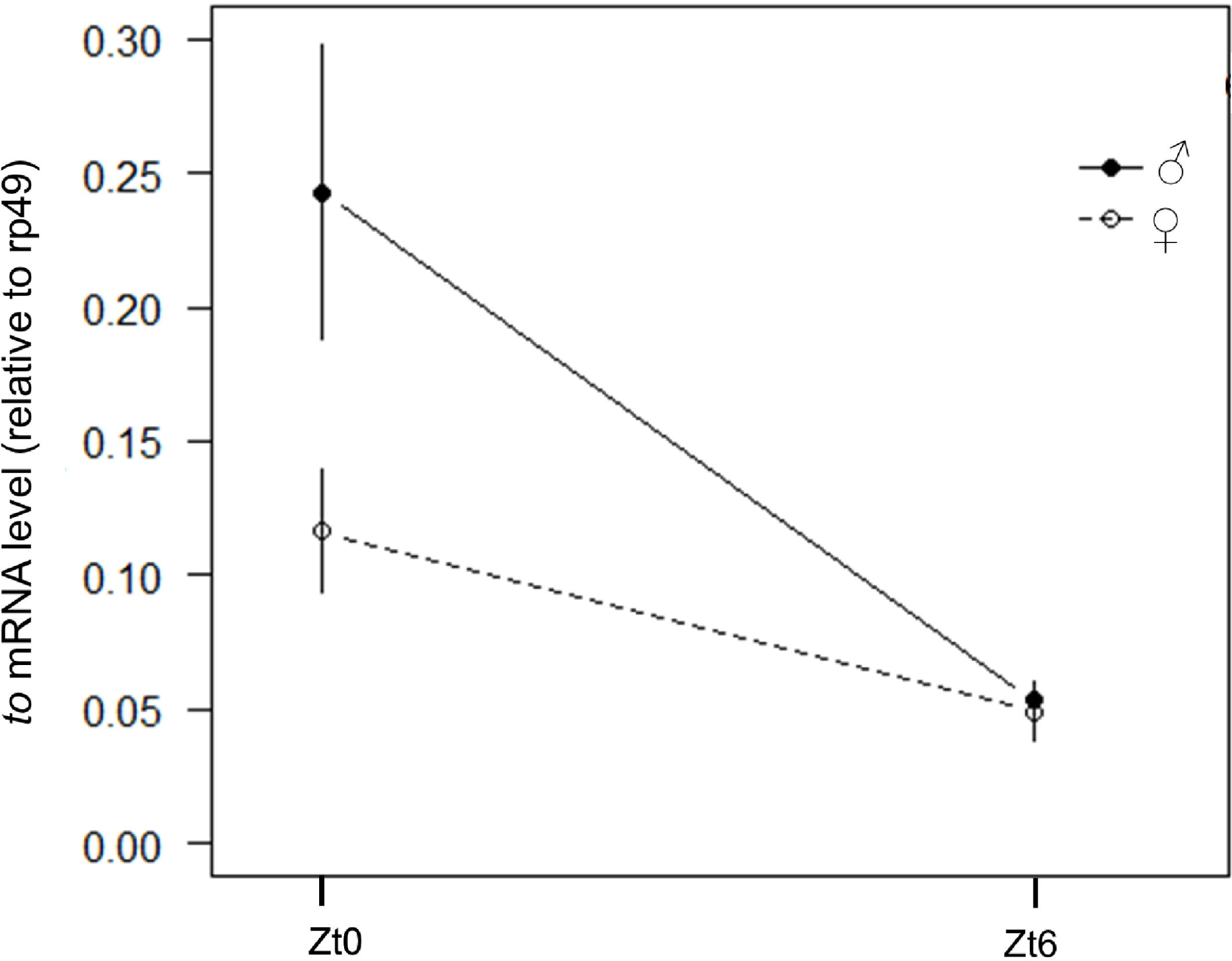
Sexual dimorphism in *takeout* expression. The relative mRNA expression of males (filled circles) and females (open circles) is depicted for Zt0 and Zt6. Expression is normalised to reference gene *rp49*. The error bars represent SE.

The sexual dimorphism in sleep was also attributed to the egg-laying activity of females (2010), which in flies is also under circadian-clock regulation (Sheeba et al., 2001). Oviposition by itself, cannot explain the reduced mid-day sleep, since it peaks after dusk (Sheeba et al., 2001), but females may need to be active during mid-day for acquiring nutrients for egg production, and these gender-specific metabolic constraints may underlying the sleep sexual dimorphism. However, in the current study only young virgin females have been used, so this excludes oviposition being a major factor for lack of siesta in females that we have observed (in all GAL4 and UAS strains, as well as Canton-S). This is also in apparent contradiction to Isaac et al. (Isaac et al., 2010) who reported that virgin females show male-like siesta, and switch to mid-day activity following mating because of the effect of the sex-peptides (SP) transferred by the males. However, the substantial lower day sleep in virgin females compared to males that we observed was also reported by others (Harbison et al., 2009). The discrepancy between the studies may be due to the different strains used, but in general, other mechanisms in addition to the SP seem to contribute to the decreased mid-day sleep of females. These mechanisms may include both neural and non-neural circuits as suggested by the current work.

## Acknowledgment

We thank CP Kyriacou and E Rosato for comments and suggestions that helped improve and clarify this manuscript and Mirko Pegoraro for his technical guidance. We are grateful to Brigitte Dauwalder, Jean-François Ferveur for sharing fly strains. This work was funded by grants from BBSRC BB/G02085X/1 to ET.

